# Integrated proteomic screening reveals design principles of CRBN molecular glue degraders

**DOI:** 10.64898/2026.03.08.710269

**Authors:** Bachuki Shashikadze, Ines Scheller, Denise Winkler, Patrick R.A. Zanon, Anastasia Bednarz, Denis Bartoschek, Sophie Machata, Tobias Graef, Uli Ohmayer, Bjoern Schwalb, Henrik Daub, Martin Steger

**Affiliations:** NEOsphere Biotechnologies GmbH, Fraunhofer Str. 1, Martinsried, Germany

## Abstract

Cereblon (CRBN)-based molecular glue degraders (MGDs) induce the degradation of diverse disease-relevant proteins, underscoring their broad therapeutic potential. Here we systematically expand the CRBN neosubstrate landscape using a target-agnostic discovery approach. By integrating deep proteomic and ubiquitinomic profiling of a 960-compound library, we identify compound-induced ubiquitination and depletion of over 230 endogenous proteins. Among these, 124 represent previously unreported CRBN neosubstrates, with over half lacking a predicted G-loop degron. We provide this dataset via an interactive resource, [NeosubstratesDB](https://neoverse-neosubstratesdb.share.connect.posit.cloud/). Complementary cellular and biochemical assays mechanistically define the interaction domain of IRAK1 and establish G-loop-dependent degradation for BCL6. Interpretable machine learning (iML) integrating proteomic profiles with chemical structures highlights key molecular fingerprints driving neosubstrate selectivity for targets such as CSNK1A1, ZFP91 and WEE1. Together, these findings significantly expand the repertoire of CRBN neosubstrates and provide a framework for rational design of next-generation MGDs.

## Introduction

Molecular glue degraders (MGDs) are small molecules that selectively reprogram the surface of E3 ubiquitin ligases to recruit and induce proteasomal degradation of non-native substrates, termed neosubstrates^1^. Cereblon (CRBN), a substrate receptor of the Cullin-Ring ligase 4 (CRL4^CRBN^) complex, has proven to be an exceptionally tractable ligase for targeted protein degradation (TPD). Indeed, the first CRBN-based MGD thalidomide and its derivatives lenalidomide and pomalidomide have become a cornerstone of multiple myeloma therapy^2–5^. Systematic derivatization of these immunomodulatory drugs (IMiDs) has yielded numerous MGDs with distinct neosubstrate selectivities, possessing a characteristically steep structure-activity relationship (SAR) where minor chemical modifications can profoundly shift neosubstrate degradation profiles^6–9^. While rational design principles for well-characterized CRBN neosubstrates are beginning to emerge, the systematic mapping of chemical structures to proteome-wide degradation remains a critical bottleneck. Expansive and publicly accessible datasets that correlate these structural motifs with degradation profiles of selected neosubstrates are indispensable to fuel the development of predictive models for TPD.

Early structural studies of CRBN-MGD-neosubstrate ternary complexes identified a characteristic three-dimensional β-hairpin α-turn (termed the G-loop) containing an invariant glycine residue that mediates interaction with MGD-engaged CRBN^10,11^. Proteome-wide computational mining predicted more than 1,600 CRBN-compatible G-loop-containing proteins, roughly 650 of which harbor a C_2_H_2_ zinc finger domain^12^. Subsequent studies uncovered MGD-induced degradation of proteins lacking a G-loop, suggesting that the neosubstrate space accessible to CRBN-based degraders extends beyond G-loop motifs^12–14^. Various zinc finger domain types have since been isolated and systematically evaluated for their susceptibility to MGD-induced engagement or degradation using reporter-based in vitro interaction assays and cellular degradation readouts^15,16^. Protein complementation assays performed in yeast with isolated protein domains independently validated many of these findings and further identified several non-G-loop-containing neosubstrates as CRBN binders through a combination of surface mimicry and complementation-based screening^17,18^. Collectively, these studies have characterized dozens of proteins as CRBN binders using isolated domain screening and a limited set of tool compounds, but it remains unclear how readily such ‘silent’ CRBN binders can be converted into productive degradation events.

Mass spectrometry (MS)–based affinity enrichment (AE-MS) with purified CRBN as bait has significantly expanded the neosubstrate landscape, identifying hundreds of compound-dependent interactors^19–21^. However, these datasets are frequently confounded by the co-purification of large protein complexes, requiring extensive downstream validation to distinguish direct from indirect binders. While global proteomics remains the gold standard for unbiased, systematic detection of degrader-induced protein depletion^6,22,23^, it cannot, on its own, discriminate between direct and indirect degrader effects. Moreover, large scale, publicly accessible proteomics datasets of CRBN-based MGDs that could be leveraged for machine-learning-guided compound design strategies remain scarce. Consequently, our understanding of the endogenous protein targets of CRBN-based MGDs, as well as the precise molecular determinants governing their selectivity, remains incomplete.

Here, we address these gaps using an integrated, target-agnostic screening strategy that combines deep proteomic and ubiquitinomic profiling with chemoinformatic analyses. By systematically interrogating a large library of CRBN-based MGDs, we substantially expand the accessible neosubstrate landscape. We identify active degraders for both novel neosubstrates and numerous previously reported CRBN binders and derive design principles that guide the rational development of next-generation MGDs. These screening results are provided as a resource to the scientific community via an interactive database ([NeosubstratesDB](https://neoverse-neosubstratesdb.share.connect.posit.cloud/)).

## Results

### In-depth characterization of a molecular glue library by proteomics

Building on our previous work that integrated proteomics screening is a powerful tool for unbiased neosubstrates discovery at scale, we sought to determine if the CRBN neosubstrate landscape could be further expanded by increasing chemical diversity^6^. While our previous work was limited to 100 molecules, we here scaled this analysis by nearly an order of magnitude. Using an established 96-well proteomics screening workflow coupled with data-independent acquisition mass spectrometry (DIA-MS)^6^, we systematically profiled 960 CRBN-based MGDs. We selected HEK293 cells overexpressing CRBN as our screening model, as this system provides enhanced sensitivity for capturing MGD-induced protein regulation^6^. To balance efficient neosubstrate discovery with compound-induced secondary downstream effects that might obscure direct degrader effects, we selected a 6-hour treatment duration. To maximize screening capacity, we optimized our previously established workflow by transitioning from triplicate to duplicate measurements. This modification allowed us to accommodate 40 compounds per plate (compared to 27 in our previous study^6^), effectively increasing the throughput by about 50% without compromising data quality (Fig. 1a). On average, we quantified over 10,000 protein groups per 96-well plate, yielding a comprehensive data matrix of more than 20 million protein quantifications with high quantitative precision (median coefficients of variation of ∼7%) (Supplementary Fig. 1a,b and Supplementary Data 1). Seventeen compounds were excluded from further analyses due to high inter-replicate variance, which resulted in apparent false-positive protein modulations. We classified the remaining 943 compounds according to their activity, defined as the total number of statistically significant protein downregulations (Fig. 1b). While approximately 27% of molecules were inactive, 526 (∼56%) downregulated ten or fewer proteins, representing promising hits with direct neosubstrate degradation activity. In contrast, 57 compounds (∼6%) displayed high activity, defined as downregulation of more than 100 proteins. For the majority of these, GSPT1 or GSPT2 protein levels were reduced by more than 75%, and the magnitude of GSPT1 fold-decrease strongly correlated with the total number of downmodulated proteins, consistent with secondary effects resulting from impaired protein synthesis (Supplementary Fig. 2). Finally, 107 degraders induced downregulation of more than 10 and up to 100 proteins and were therefore classified as medium-active compounds.

**Figure 1.**
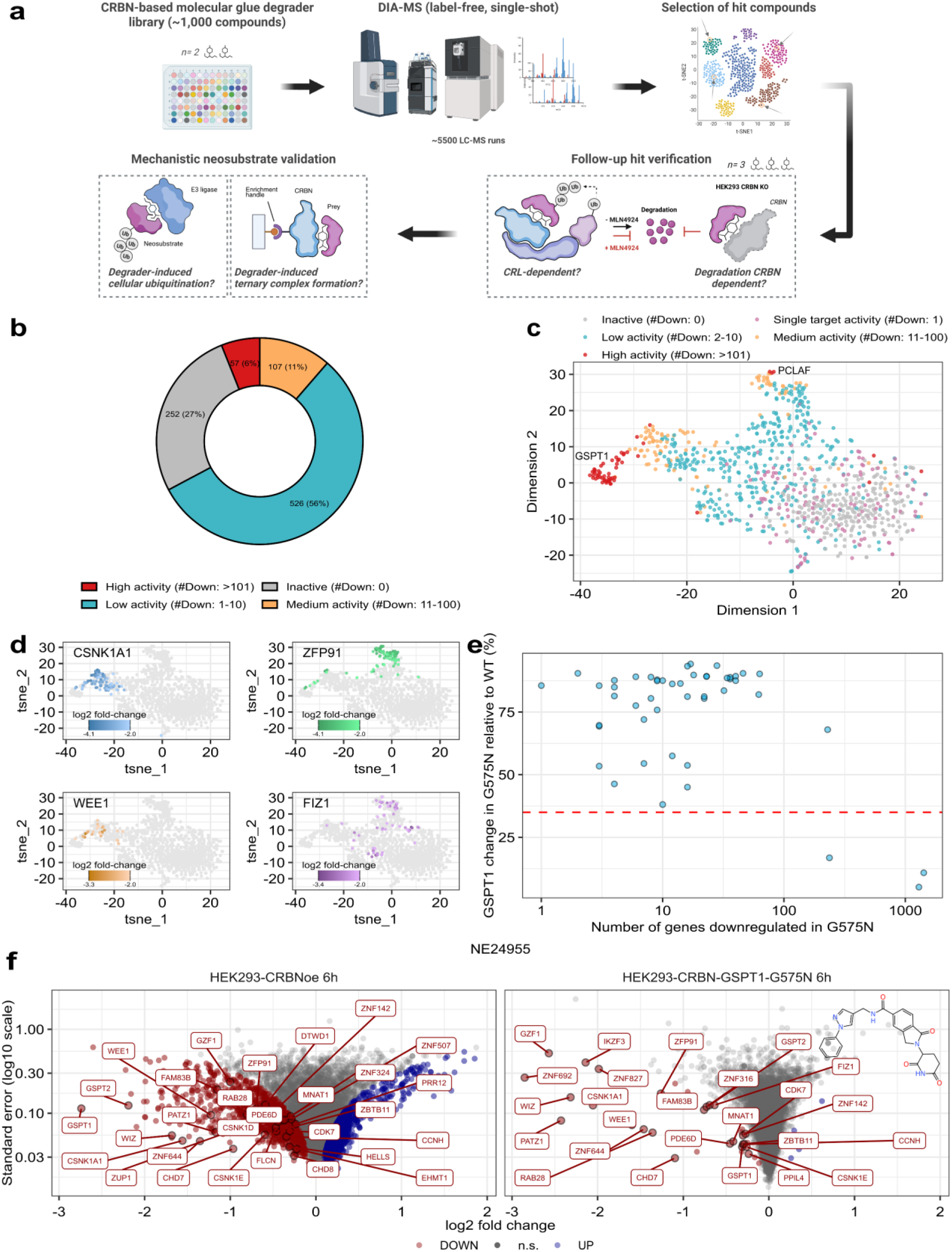
Proteomics screening of a 960 molecular glue degrader (MGD) library and compound classification. **a** Schematic of the proteomics screening workflow. HEK293 cells overexpressing CRBN were cultivated in 96-well plates and treated with compounds for 6 hours (duplicates per compound). Proteins were extracted, digested, and the resulting peptides analyzed by label-free data-independent acquisition mass spectrometry (DIA-MS). The data were reviewed and hit compounds selected for follow-up verification, which involved retesting in triplicate with or without MLN4924 to assess cullin–RING ligase (CRL) dependency. CRBN dependency was evaluated in HEK293-CRBN⁻^/^⁻ cells. Neosubstrate candidates were mechanistically validated by global ubiquitinomics, and ternary complex formation was assessed for selected candidates. **b** Classification of compounds according to their activity, based on the number of significantly (adj. p-value < 0.01) downregulated proteins. **c** t-SNE visualization of proteome profiles across all tested compounds. Activity classes are highlighted with different colors. GSPT1 and PCLAF clusters are highlighted. **d** Same t-SNE visualization as in **c**, with measured log_2_ fold-changes (< -2) of selected neosubstrates overlaid onto the plot. **e** Scatter plot showing number of significantly (adj. p-value < 0.01) downregulated proteins (x-axis) and ratio of GSPT1 log_2_ fold-change in HEK293 cells overexpressing CRBN relative to cells co-expressing CRBN and the degradation resistant mutant GSPT1-G575N (y-axis). The dashed red horizontal line indicates the 35% threshold, below which GSPT1-dependent secondary effects become evident. **f** Volcano plots showing log_2_ fold-change (x-axis) versus log_10_ standard error (y-axis) for all quantified proteins in HEK293 cells overexpressing CRBN, either alone (left) or in combination with GSPT1-G575N (right), following 6-hour treatment with NE24955. Statistically significant (adj. p-value < 0.01) up- and downregulated proteins are colored in blue and red, respectively. n.s.= not significant. Selected proteins are labeled.

To gain deeper insight into compound activity and the underlying protein regulatory landscape, we visualized the proteomic signatures of the curated set (943 compounds) using t-SNE analysis. This revealed a distribution along an activity gradient, with high activity compounds partitioning into one large and one smaller distinct cluster, suggesting shared protein modulation patterns. Within these clusters, the most strongly downregulated proteins were either the neosubstrate GSPT1 or PCLAF, a PCNA-associated factor which we previously identified as a common off-target of CRBN MGDs^6,10^ (Fig. 1c). To further explore regulation patterns, we mapped fold-changes of several frequently downregulated neosubstrates (CSNK1A1, WEE1, ZFP91 and FIZ1) onto the t-SNE space (Fig. 1d). This uncovered a prominent subset of CSNK1A1 degraders co-targeting WEE1. In contrast, ZFP91 degraders clearly segregated from CSNK1A1-targeting molecules, indicating highly divergent downstream protein modulation profiles. FIZ1 degraders were generally more selective than those targeting CSNK1A1 or WIZ, yet they displayed a more scattered distribution across the t-SNE space, suggesting the co-regulation of additional neosubstrate unique to each sub-cluster. While low-activity compounds generally did not form clear clusters, we identified a distinct subset of 18 compounds that segregated sharply from the rest (Supplementary Fig. 3a). Examination of their protein regulation profiles showed consistent CRBN downregulation, and a closer inspection of the data revealed two classes of degraders: one showing selective CRBN depletion, likely driven by ligand-induced homodimerization as previously described^24^, and another comprising molecules with CRBN/DDB1 degrader activity. Notably, this latter class was characterized by the coregulation of the substrate receptor TRPC4AP, suggesting targeted degradation of the CRL4^TRPC4AP^ complex (Supplementary Fig. 3b,c).

To unmask putative neosubstrates that might otherwise be obscured by secondary regulation, we re-screened 50 potent GSPT1 degraders in a HEK293 line co-expressing CRBN and the G-loop degron mutant GSTP1-G575N^10^ (HEK293_GSPT1-GN) (Supplementary Fig. 4). We reasoned that the overexpression of this non-degradable mutant would compensate for the loss of endogenous GSPT1 and thus preserve physiological protein translation while uncovering direct CRBN neosubstrates other than GSPT1. For 47 of the tested compounds, GSPT1 depletion was indeed decreased by more than 35% in the HEK293_GSPT1-GN line, which was sufficient to effectively suppress GSPT1-driven secondary effects. By mitigating these translation defects, we successfully unmasked numerous previously obscured neosubstrates and neosubstrate candidates that would have otherwise remained undetected in the wild-type background (Fig. 1e,f). Of note, three compounds induced robust GSPT1 degradation despite GSPT1-G575N overexpression, indicating that their high intrinsic degradation potency can compensate for the G-loop mutation, consistent with prior observations for highly potent SALL4 degraders^16^ (Supplementary Fig. 5).

To prioritize lead molecules and their associated neosubstrate candidates for follow-up validation, we applied a two-fold strategy. First, we systematically reviewed significant protein downregulations induced by low- and medium-active degraders, including GSPT1-targeting compounds analyzed in the HEK293_GSPT1-GN rescue line (Methods). This process resulted in the nomination of 72 neosubstrate candidates from the primary screen and 13 additional candidates uniquely identified in the HEK293_GSPT1-GN line (Supplementary Data 2). Second, we incorporated several highly promiscuous, low-selectivity degraders into our validation set. These molecules were selected for their ability to simultaneously degrade dozens of diverse neosubstrates, similar to our previously identified MGD SJ42101^6^. Finally, we added several positive controls regulating well characterized neosubstrates.

To increase confidence in the identified neosubstrate candidates and assess their cullin-RING ligase (CRL) dependency, we retested representative hit compounds for each candidate in a 6-hour treatment experiment, both with and without the neddylation inhibitor MLN4924^25^.

GSPT1 degraders were re-tested in HEK293_GSPT1-GN cells (Supplementary Data 3). Most candidates showed complete or near-complete rescue of regulation upon MLN4924 treatment, confirming their dependence on CRL activity. Conversely, we identified a subset of CRL-independent proteins, including the known MLN4924-insensitive factor PCLAF^6^, as well as TXNIP, HMGCR and WLS (Supplementary Fig. 6). Collectively, our proteomic screen of 960 compounds enabled a systematic classification by activity, clustering by shared regulation patterns, and the identification of numerous MLN4924-sensitive neosubstrate candidates, including many uniquely uncovered via the degradation-resistant HEK293_GSPT1-GN line.

### Large-scale validation of candidate neosubstrates

Our screen identified hundreds of proteins downregulated in a MLN4924-dependent manner, nominating them as putative CRBN neosubstrates. To distinguish direct targets from secondary downregulation events, we conducted a series of ubiquitinomics experiments using a representative set of 83 compounds. This selection included at least one hit for each MLN-sensitive candidate, all identified promiscuous degraders and several positive controls. For global ubiquitination site mapping, we employed di-Gly (or K-GG) remnant peptide profiling coupled with slice-PASEF^26–28^. This technology utilizes tryptic digestion of ubiquitinated proteins, to generate peptides carrying a characteristic di-Glycine remnant on modified lysine residues. Enrichment and MS-based analysis of these peptides enable site-specific mapping and quantification of ubiquitination events with high sensitivity on a proteome-wide scale. Given that primary intracellular neosubstrate ubiquitination occurs within minutes after MGD exposure, we employed a 30-minute treatment window to capture ubiquitination events prior to detectable changes in total protein levels. Ubiquitination within this short timeframe is considered highly indicative of a direct, proximity-induced relationship between the neosubstrate and the CRBN-bound MGD^6,29^.

In total, we quantified 69,434 ubiquitination sites mapping to 9,926 proteins. 186 of those sites were detected exclusively following compound treatment and were absent under basal conditions, suggesting they occur independently of physiological protein turnover. To further validate this observation, we cross-referenced these selectively induced sites against two large-scale, publicly available ubiquitination site datasets, collectively comprising more than 100,000 sites^26,30^. Notably, 85% of compound-induced sites were absent from these datasets, reinforcing their classification as de novo ‘neo-ubiquitination’ events that occur exclusively through induced proximity to CRBN-MGD in the context of targeted protein degradation (Supplementary Fig. 7).

Many CRBN neosubstrates are engaged and ubiquitinated by promiscuous MGDs, yet not to an extent sufficient to trigger proteasomal degradation. Numerous such ‘latent’ neosubstrates have previously been identified using MS-based interactomics, ubiquitinomics, and protein complementation assays^6,12,17,19–21^. Furthermore, while reporter-based degradation screens using isolated domains have nominated a vast array of zinc-finger proteins as putative targets, the endogenous degradation of the corresponding full-length proteins has yet to be demonstrated for most of these candidates^16^.

To assess if MGD-induced ubiquitination leads to productive protein depletion, we overlaid MS-quantified protein abundance changes (6-hour treatment) with our global ubiquitinomics data (30-minute treatment). This integrative approach allowed us to dissect productive degradation events, characterized by concurrent protein ubiquitination and depletion, from cases of ubiquitination without a detectable decrease in abundance. In total, we identified 233 high-confidence candidates exhibiting both MGD-induced ubiquitination and significant protein depletion (Supplementary Data 4).

To validate the robustness and the sensitivity of our platform, we comprehensively reviewed these candidates for prior experimental evidence in the literature. For 67 of these proteins, MGD-dependent intracellular protein depletion had been previously reported (Supplementary Fig. 8). An additional 42 proteins had been nominated as putative CRBN neosubstrates based on in vitro or in vivo ternary complex assays, ubiquitination studies, or zinc-finger reporter degradation screens; however, no active degraders of the endogenous proteins have been reported to date (Fig. 2a and Supplementary Data 4).

**Figure 2.**
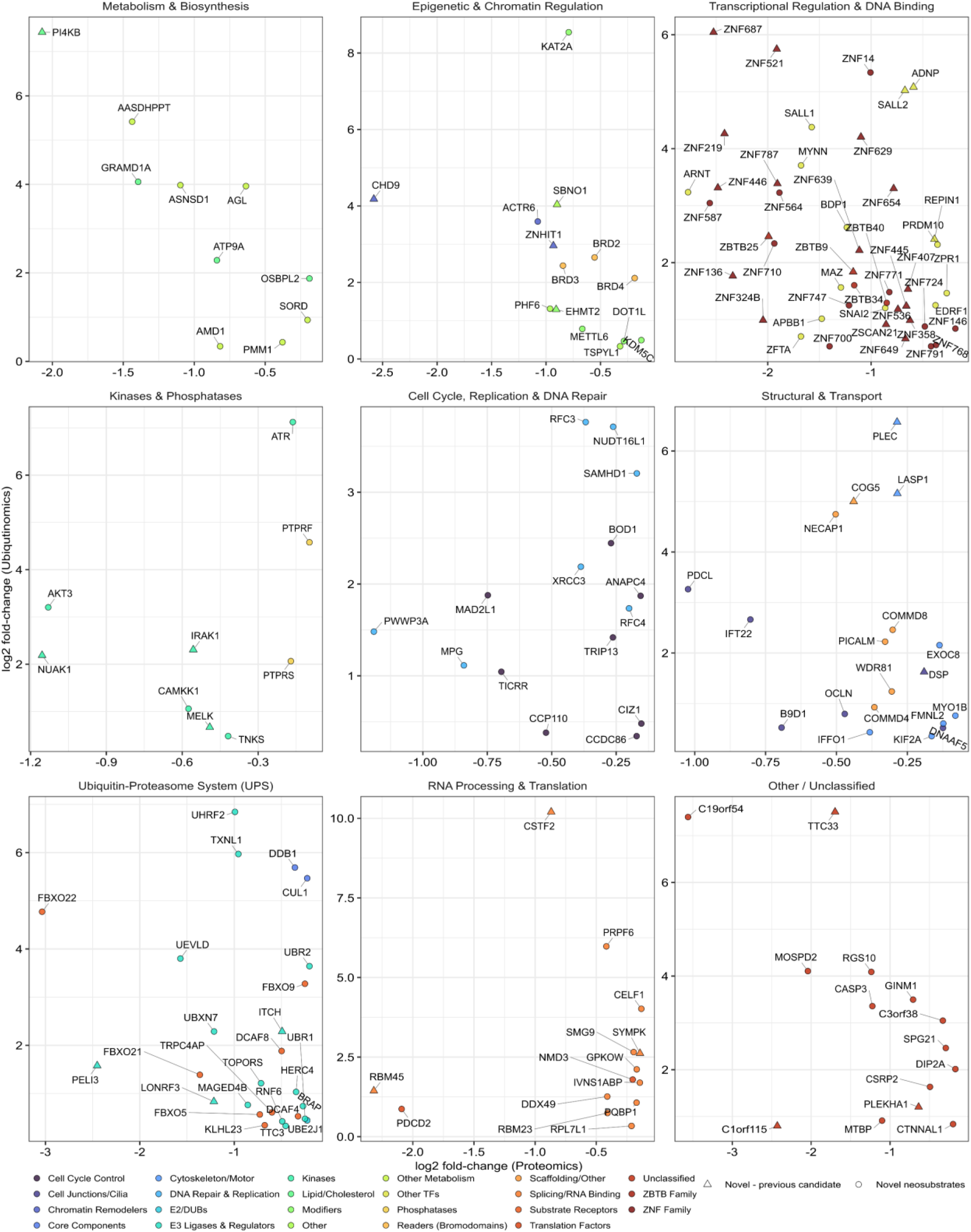
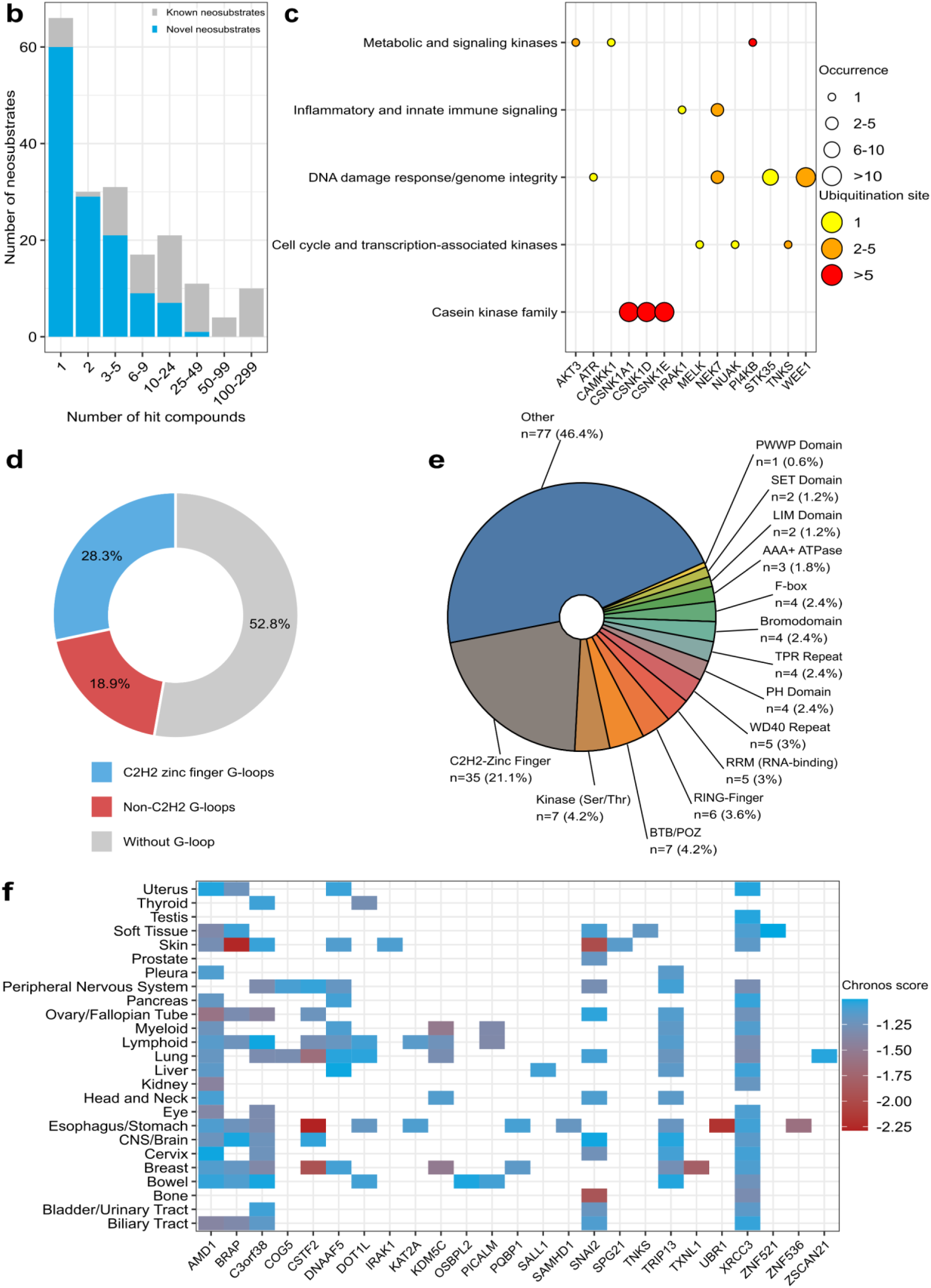
Comprehensive characterization of the CRBN neosubstrate landscape reveals functional diversity. **a** Scatter plot of proteomics log₂ fold-change (x-axis) versus ubiquitinomics log₂ fold-change (y-axis) for novel neosubstrates (circles) and previously reported candidates (triangles). For each protein, the largest detected log₂ fold decrease is plotted on the x-axis. The y-axis shows the log₂ fold induction of the most strongly upregulated ubiquitination site induced by the same compound. The neosubstrates are grouped according to molecular function. See [NeosubstratesDB](https://neoverse-neosubstratesdb.share.connect.posit.cloud/) for further details. **b** Bar chart showing the number of degrader compounds (x-axis) versus the number of neosubstrate occurrences (y-axis). Neosubstrates are further classified as previously reported (grey) or newly identified in this study (blue). **c** Representation of kinases identified as CRBN neosubstrates in the proteomic screen, grouped by kinase family. Dot size reflects the number of occurrences of significant downregulation (adj. p-value < 0.01), while color indicates the number of ubiquitination sites significantly induced (adj. p-value < 0.05) upon compound treatment. **d** Pie chart categorizing proteins according to the presence of a predicted G-loop degron. G–containing neosubstrates were further classified according to G-loop type (C_2_H_2_ zinc finger and non-C_2_H_2_)^12^. **e** Pie chart showing distribution of neosubstrates across selected structural domain classes. **f** Gene essentiality profiles based on DepMap^35^ cancer dependency scores. For each category, chronos scores < −1 were averaged and reported as a combined score.

The remaining 127 proteins represent novel candidates not previously described in the literature. To our surprise, ubiquitinomics revealed clear, compound-induced ubiquitination for four MLN4924-insensitive targets (INPP5E, RAB28, WLS and HMGCR). Retesting all hit compounds in CRBN^−/−^ cells confirmed their CRBN-independent regulation (Supplementary Fig. 9). While RAB28 was regulated in a CRBN-dependent manner by over 20 compounds, its specific co-regulation with the ciliary phosphatase INPP5E by one compound was CRBN-independent, suggesting a localized off-target effect. Similarly, the CRBN-independent ubiquitination and depletion of WLS or HMGCR likely reflect compound-induced ER stress rather than direct CRBN recruitment. Accounting for these off-targets, we identified 124 high-confidence, novel CRBN-dependent neosubstrate candidates (Fig. 2a). Details on neosubstrate regulation across different cellular context and treatment times, as well as information on compound-induced ubiquitination sites can be accessed at [NeosubstratesDB](https://neoverse-neosubstratesdb.share.connect.posit.cloud/).

Our compound library consisted primarily of IMiDs, defined by a conserved glutarimide ring linked to a variable phthalimide or isoindolinone moiety^31^. Consequently, well-described IMiD neosubstrates such as CSNK1A1, CSNK1E, FIZ1, GSPT1, GZF1, WIZ and ZFP91 were among the most frequent screening hits (Supplementary Fig. 10a). Detailed inspection of the screening data further revealed a broad array of highly selective and efficacious degraders for several previously reported neosubstrates, such as WBP4, PRR12 and CNOT4^6,12^ (Supplementary Fig.10b). Conversely, 60 neosubstrates were targeted by only one or two compounds, including several selectively regulated disease-relevant proteins. Among these were CASP3, a known driver of early pathogenesis in Alzheimer’s disease^32,33^, and AKT3, a kinase for which PROTAC-mediated degradation has previously demonstrated *in vivo* antitumor efficacy in non-small cell lung cancer models^34^ (Supplementary Fig. 11). Additionally, we uncovered degraders targeting multiple other kinases, including the DNA damage associated serine/threonine-protein kinase ATR (ATR), calcium/calmodulin-dependent protein kinase kinase 1 (CAMKK1), Interleukin-1 receptor-associated kinase 1 (IRAK1), phosphatidylinositol 4-kinase beta (PI4KB), and maternal embryonic leucine zipper kinase (MELK) (Fig. 2b,c). Beyond kinases, the identified neosubstrates spanned multiple classes of transcriptional regulation, including sequence-specific DNA-binding transcription factors (*e.g.*, BCL6, ARNT), more than 40 C2H2 zinc-finger transcription factors (*e.g.* ZBTB and ZNF family members), chromatin remodelers (*e.g.* CHD7/8/9, HELLS) and epigenetic and transcriptional co-regulators (*e.g.* DOT1L, MYNN) (Supplementary Fig. 12).

About 53% of the identified neosubstrates lacked a predicted G-loop degron, including the previously reported, G-loop-independent target G3BP2^12,13^. Within this subset, roughly 30% harbored a C_2_H_2_ zinc finger domain (Fig. 2d and Supplementary Data 4).

Representative protein domains other than C_2_H_2_ zinc fingers included Bromodomains, BTB/POZ, RING-finger and RNA-binding domains (Fig. 2e). Assessment of clinical relevance revealed that 15% of the identified neosubstrates are classified as strongly selective dependencies according to DepMap^35^. Furthermore, cross-referencing our candidates with the Open Targets^36^ platform uncovered robust genetic and pathological associations spanning a broad range of non-oncological diseases (Fig. 2f and Supplementary Fig.13).

Neosubstrate recruitment to CRBN can either occur directly or indirectly. Indirect neosubstrates (known as bystander or piggyback neosubstrates) are components of protein complexes that are recruited to CRBN-MGD complexes via their physical association with primary neosubstrates^6,12,13^. To distinguish direct neosubstrates from potential bystanders, we leveraged established protein-protein interaction (PPI) networks for our list of neosubstrates. We assessed the co-occurrence of downregulation for identified protein pairs across the compound library and integrated physical association confidence scores from STRING^37^ with normalized co-occurrence frequencies (i.e., protein pairs depleted together more often than in isolation). This approach identified several cases in which loss of a neosubstrate is likely to reflect degradation of a physically associated primary target. Examples include WIZ-ZNF644-EHMT2 (with WIZ as the primary target), as well as CRBN-DDB1-TRPCAP4, ACTR6-ZNHIT1 and FBXO5-CDC20 (Supplementary Fig. 14).

Finally, we assessed whether the depletion of our identified neosubstrates could be recapitulated in cells with native CRBN expression. To this end, we measured the regulation of a representative panel of candidates in parental HEK293 cells following treatment with their respective hit compounds for 6 and 24 hours. This analysis confirmed that 47% of the tested candidates were significantly depleted at one or both time points in parental HEK293 cells, demonstrating high translatability of our screening hits into a physiological cellular context (Supplementary Fig. 15).

### Mechanistic insights into IRAK1 and BCL6 neosubstrate recognition

Our screen revealed an approximately even distribution of neosubstrates with and without a predicted G-loop^12^, including numerous disease-relevant targets. We therefore selected one representative neosubstrate from each class for mechanistic characterization. IRAK1 is a central node in the Toll-like receptor and interleukin signaling pathways, possessing both kinase activity and scaffolding functions^38^. The latter has been implicated in inflammation and chemoresistance, supporting IRAK1 as an attractive candidate for degrader-based therapeutic intervention^39,40^. Prior structural modeling failed to identify a G-loop degron, and prior MS-based affinity enrichment mass spectrometry (AE-MS) studies categorized it as CRBN-pomalidomide binder without evidence of degradation^19^. Here, we demonstrate ubiquitination at K239, K355 and K397, and intracellular depletion of IRAK1 induced by NE24878 in both parental HEK293 cells and CRBN-overexpressing cells, with selectivity over IRAK4 (Fig. 3a). As a proxy for a disease-relevant cell type, we examined whether IRAK1 degradation could be recapitulated in human peripheral blood mononuclear cells (PBMCs). Treatment of freshly isolated PBMCs with NE24878 for 24 h resulted in a selective, about two-fold reduction of IRAK1 levels with clear selectivity over other IRAK family members (Fig. 3a). IRAK1 is composed of an N-terminal death domain (DD), a central kinase domain, and a C-terminal domain (CTD) containing TRAF6 binding motifs. Structurally, the kinase domain is organized into N- and C-lobes which harbor the ATP-binding pocket and catalytic loop, respectively. Within the IRAK family, the DD domain and kinase N-lobe are well conserved, while the C-lobes and the C-termini exhibit high divergence. We set out to map the specific domain mediating IRAK1-CRBN interaction, hypothesizing that these less conserved regions were more likely to mediate the binding. Using AE-MS with as a read-out, we assessed compound-meditated in vitro interaction between recombinant CRBN^midi^ and IRAK1 truncation mutants spanning residues 1–524 (lacking the CTD) and 1–381 (lacking both the CTD and the kinase C-lobe)^41^. Both endogenous IRAK1 and the 1-524 fragment showed specific CRBN interaction, whereas overexpressed full-length IRAK1 did not, likely due to aggregation mediated by the unstructured CTD. In contrast, the 1-381 fragment showed no detectable interaction, pinpointing residues 381-524 as critical for interaction with CRBN (Fig 3b).

**Figure 3.**
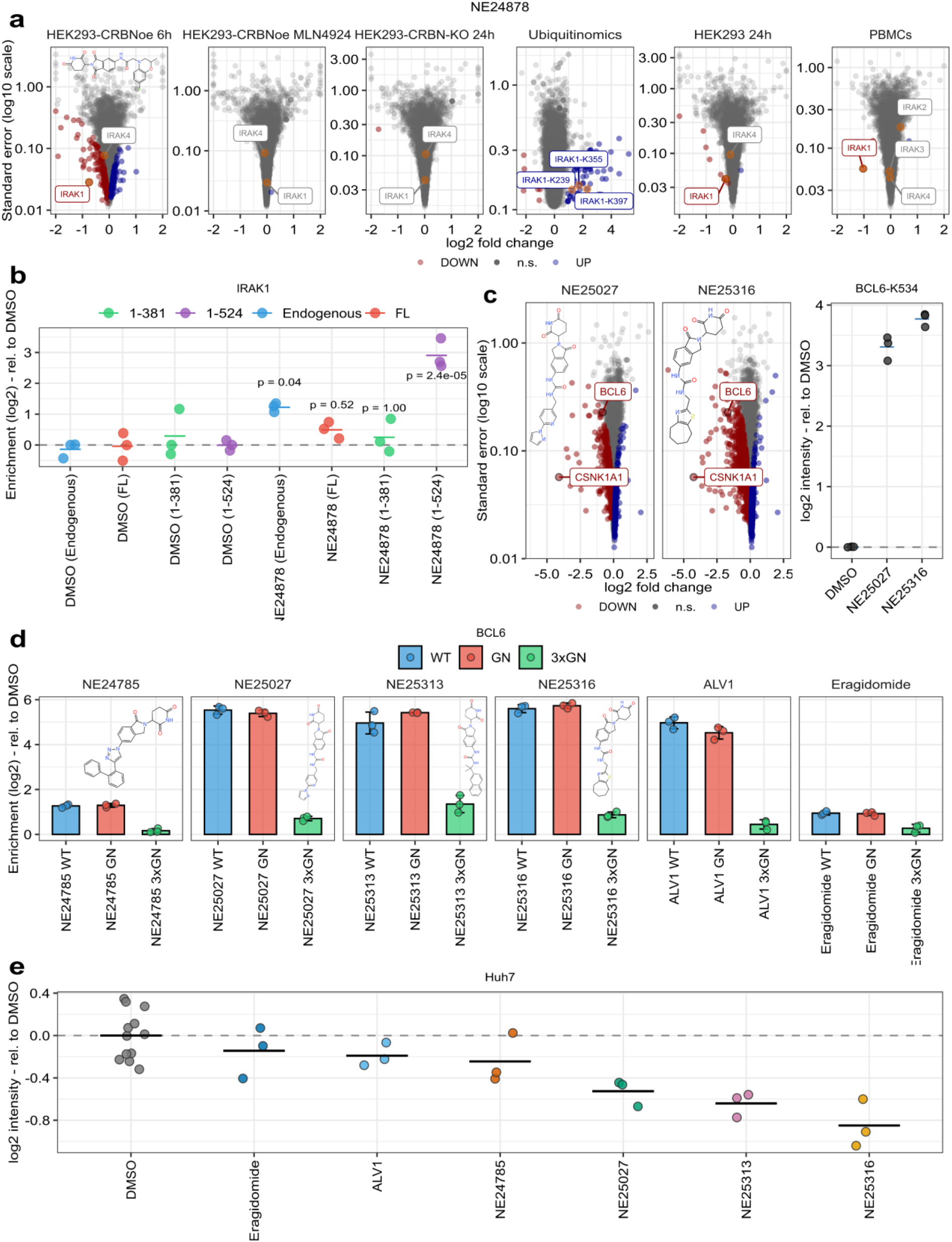
Characterization of IRAK1 and BCL6 as CRBN neosubstrates. **a** Volcano plots displaying log₂ fold change (x-axis) versus −log₁₀ adjusted p-value (y-axis) for proteins or ubiquitination sites following treatment with NE24878. From left to right, the plots show: (i) compound-treated HEK293_CRBNoe cells (6 hours) in the absence of MLN4924, (ii) co-treatment with MLN4924, (iii) protein regulation in HEK293 CRBN⁻/⁻ cells (24 hours), (iv) quantified ubiquitination sites after a 30-minute compound treatment (v) parental HEK293 (24 hours) and (vi), Peripheral blood mononuclear cells (PBMCs) treated for 6 hours. Statistically significant up- and downregulated proteins/sites (adjusted p-value < 0.01 for proteomics and < 0.05 for ubiquitinomics) are highlighted in blue and red, respectively; n.s.= not significant. Selected proteins/sites are annotated. **b** IRAK1 protein quantification by affinity enrichment mass spectrometry (AE-MS). Full-length IRAK1 (1–712) and truncation constructs (1–381 and 1–524) were transiently expressed in HEK293 cells. Lysates were treated with 10 µM NE24878 in the presence of biotinylated CRBN^midi^, followed by streptavidin-mediated bait enrichment and MS-based quantification. A non-transfected control was included to enable quantification of endogenous IRAK1. The x-axis indicates enrichment relative to DMSO control. Reported p-values were adjusted for multiple hypothesis testing using the Benjamini–Hochberg method. **c** Volcano plots displaying log₂ fold change (x-axis) versus −log₁₀ adjusted p-value (y-axis) for proteins quantified in HEK293-CRBNoe cells treated for 6 hours with NE25027 (left) or NE25316 (right). Statistically significant up- and downregulated proteins (adjusted p-value < 0.01) are highlighted in blue and red, respectively; n.s., not significant. Selected proteins are annotated. The dot plot on the right shows quantification of the BCL6-K534 ubiquitination site in HEK293-CRBNoe cells treated with the respective compounds for 30 minutes. **d** BCL6 protein quantification by affinity enrichment mass spectrometry (AE-MS). Wild-type (wt), G608N (GN) or G598N/G608N/G636N (3xGN) BCL6 constructs were transiently expressed in HEK293 cells. Lysates were treated with 10 µM of the indicated compounds in the presence of biotinylated CRBN^midi^, followed by streptavidin-mediated bait enrichment and MS-based quantification. Erag= Eragidomide. **e** MS-based BCL6 quantification in HuH-7 cells treated with 10 µM of the indicated compounds for 6 h.

As a representative example of a disease-relevant, G-loop-containing neosubstrate, we selected B-cell Lymphoma 6 protein (BCL6). BCL6 is frequently translocated or mutated in B-cell malignancies, and elevated levels are associated with chemoresistance in both hematological and solid tumors^42–44^. While early efforts to inhibit BCL6 were hindered by its extensive scaffolding role, targeted protein degraders have either shown promise in preclinical studies or successfully entered clinical evaluation^45–48^. A reporter-based degradation screen with human zinc finger domains recently described BCL6 as a putative neosubstrate, a finding validated by the MGD-induced depletion of the endogenous, full-length protein^49^. We identified NE25316 and NE25027 as compounds significantly downregulating BCL6, although quantification precision was limited by its low expression in HEK293 cells (protein abundance rank >10,000). Both molecules also induced strong degradation of CSKN1A1, resulting in extensive secondary regulations. Notably, ubiquitinomics revealed compound-specific ubiquitination of BCL6 at K534 in response to treatment with these two molecules, corroborating direct CRBN-mediated degradation (Fig. 3c). Further inspection of our proteomics dataset identified NE25313 as an additional dual CSNK1A1 and BCL6 degrader. While compound-induced ubiquitination supported direct degradation by CRBN, we next determined whether BCL6 depletion resulted from a direct, physical interaction with CRBN or occurred as a secondary effect downstream of CSNK1A1 degradation. To address this, we analyzed compound-induced CRBN–BCL6 binding by AE-MS. As negative controls, we included a potent CSNK1A1 degrader NE24785 that did not elicit any detectable BCL6 depletion, as well as eragidomide (CC-90009)^50^, a MGD with distinct neosubstrate specificity. The recently reported BCL6 degrader ALV1^49^ served as positive control. Three putative G-loop degrons have been predicted for BCL6, centered on glycine residues G598, G608, and G636, with G608 exhibiting the lowest predicted RMSD value^12^. We therefore included a G608N (GN) mutant in the assay. Encouragingly, we detected CRBN binding of both endogenous and overexpressed wild type (wt) BCL6 in the presence of ALV1, NE25313, NE25316, and NE25027, but not NE24785 or Eragidomide (Fig. 3d). Notably, enrichment of BCL6-GN was comparable to BCL6-wt, indicating that mutation of G608 alone is insufficient to disrupt binding to the CRBN-MGD complex and suggesting that neighboring C_2_H_2_ zinc fingers may contribute to the interaction, consistent with recent observations for IKZF2^49^. We therefore generated a triple GN mutant (3×GN) by mutating all invariant glycines of the three predicted G-loops (G598, G608, G636) and included it in our AE-MS assay. This revealed a clear abrogation of binding, suggesting that G598 and/or G636, either alone or in combination with G608, coordinate the binding with CRBN-MGD complexes (Fig 3d). We further observed that compared to ALV1, BCL6 enrichment was more pronounced by NE25313, NE25316 and NE25027, suggesting higher target affinity and potentially enhanced degradation rates. To test this hypothesis, we assessed BCL6 degradation in Huh7 cells, which endogenously express significantly higher levels of BCL6 than HEK293. Consistent with our binding data, NE25313, NE25316 and NE25027 induced a substantially stronger BCL6 depletion than ALV1, demonstrating the translatability of our binding studies to cellular degradation outcomes (Fig. 3e). In summary, these results highlight how global proteomics, integrated with MS-guided biochemical validation, can readily identify novel, disease-relevant neosubstrates while pinpointing the structural determinants that facilitate MGD-induced target recruitment.

### Interpretable machine learning (iML) reveals neosubstrate-specific molecular fingerprints

The abundance profiles of over 10,000 proteins across nearly 1,000 MGDs offer a unique opportunity to identify structural features that govern CRBN-dependent neosubstrate degradation, guiding the rational design of MGDs to target specific proteins or reduce off-target effects. To this end, we employed the tree-based gradient boosting algorithm XGBoost as an interpretable machine learning (iML) approach to predict protein abundance changes from molecular fingerprints (Fig. 4a). Molecular fingerprints, which are binary vectors encoding the presence or absence of specific substructural features within a molecule, were used as input for XGBoost. The model builds an ensemble of decision trees that integrate molecular fingerprints to predict the extent of neosubstrate degradation and additionally provide feature importance scores highlighting the structural determinants of degradation. Key advantages of XGBoost include its computational efficiency and the ability to capture non-linear dependencies in sparse, high-dimensional chemical spaces, allowing high predictive accuracy. Because downregulation of many proteins by highly active compounds may often be confounded by neosubstrate competition, these compounds were excluded from the analysis.

**Figure 4.**
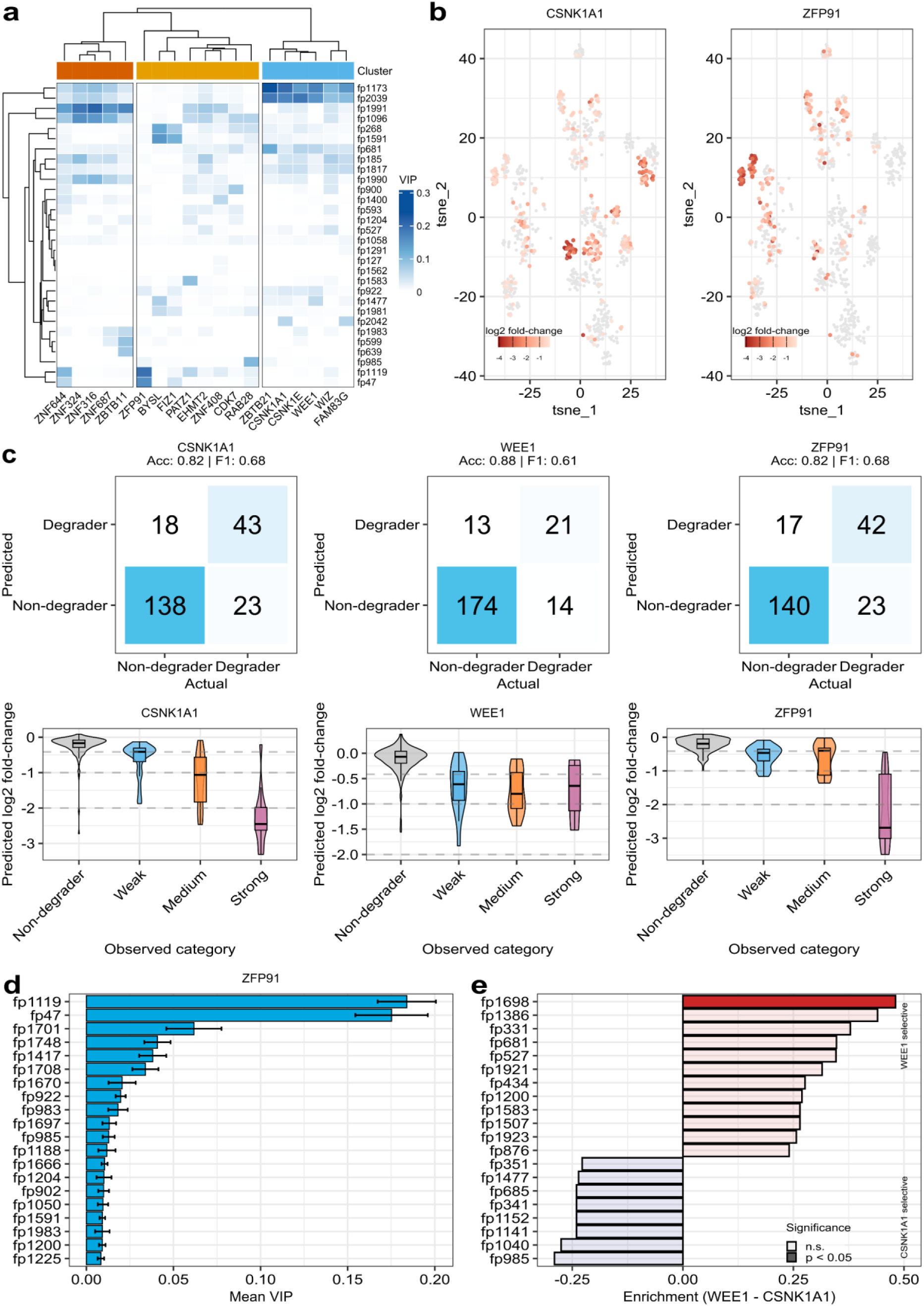
Interpretable machine learning reveals neosubstrate-specific molecular fingerprints. **a** Variable importance in projection (VIP) heatmap showing top 30 fingerprints across well-performing models (Pearson r ≥ 0.5, n = 19 targets). Hierarchical clustering reveals distinct fingerprint importance profiles for different neosubstrates, with ZFP91 and CSNK1A1 clustering separately. **b** t-SNE plot based on morgan fingerprints with measured log_2_ fold-changes of CSNK1A1 (left) and ZFP91 (right) overlaid onto the plot. **c** Confusion matrices (top) and degradation magnitude prediction (bottom) for three representative targets. Models achieved high accuracies (0.82-0.88) with low false positive rates (7-14%) and successfully distinguished weak, medium, and strong degradation categories. **d** ZFP91 variable importance plot showing top 20 fingerprints with error bars representing stability across cross-validation folds. **e** Fingerprint enrichment analysis comparing WEE1-selective versus CSNK1A1-selective degraders. Bars show enrichment scores (WEE1 - CSNK1A1) with statistical significance indicated by opacity (solid: p < 0.05, transparent: non-significant). fp1698 shows strong enrichment in WEE1-selective compounds, matching a selectivity determinant independently identified through rational medicinal chemistry.

We found that the model performance varied across the assessed neosubstrates, with Pearson correlation coefficients between predicted and observed log_2_ fold-change ranging from −0.16 to 0.82. As expected, we observed a positive correlation between model accuracy and both neosubstrate degradation frequency and degradation magnitude (Supplementary Fig. 16a,b). This likely reflects two complementary factors: first, frequently degraded proteins are represented by a larger number of positive training examples, enabling the model to learn a more robust structure–activity relationship; second, proteins degraded with high efficacy exhibit a stronger and more consistent fingerprint–degradation relationship, providing clearer structural determinants for the model to capture.

To identify structural features driving neosubstrate degradation, we analyzed the variable importance in projection (VIP) scores from our XGBoost models, which quantify each fingerprint’s contribution to overall prediction accuracy. We generated a heatmap of VIP scores for the top 30 fingerprints across neosubstrates with high-performing models (Pearson r ≥ 0.5, n = 19 targets) (Figure 4a), revealing three main clusters of neosubstrates, with each cluster defined by a distinct set of important fingerprints. This clustering highlights how specific sets of chemical features are associated with the selective degradation of different neosubstrates. Notably, ZFP91 and CSNK1A1 had the highest VIP scores and were clearly separated on the heatmap, indicating target-specific structural determinants. Consistently, t-SNE visualization of the chemical space, based on molecular fingerprints and with log₂ fold-changes highlighted for each compound, confirmed this separation, showing that the degraders of ZFP91 and CSNK1A1 occupy structurally distinct regions (Figure 4b).

Having identified target-specific structural features that distinguish MGDs, we next evaluated how well the XGBoost models could predict degradation, both as a binary classification and across different levels of degradation strength. To evaluate binary classification performance, we applied a 25% degradation threshold to assess whether the models could correctly predict compounds above or below this cutoff. Using this criterion, the WEE1, CSNK1A1, and ZFP91 models achieved accuracies of 0.88, 0.82, and 0.82, respectively, with relatively low false positive rates (7.5%, 14.3%, and 14.1%) (Fig. 4c, top panels). Additionally, the CSNK1A1 model could distinguish not only between degraders and non-degraders, but also predict degradation magnitude across weak, medium, and strong categories, whereas predictions for WEE1 and ZFP91 were less clearly stratified (Fig. 4c, bottom panels, Supplementary Fig. 17).

To understand how structural features drive degradation across multiple neosubstrates, particularly for promiscuous degraders, we analyzed fingerprint importance patterns across high-performing models (Pearson correlation > 0.8). By analyzing the top three most important features per gene, we identified both promiscuous fingerprints, shared across multiple targets, and gene-specific features driving individual neosubstrate degradation (Supplementary Fig. 18a). Co-occurrence network analysis identified clusters of promiscuous fingerprints that frequently appear together in compounds co-degrading multiple targets, suggesting structural motifs that promote simultaneous degradation. In contrast, isolated fingerprints likely drive target-specific selectivity (Supplementary Fig. 18b).

Given that promiscuous fingerprints were prevalent among multi-target degraders, we next investigated whether the wider molecular context confers further neosubstrate selectivity. To directly test this, we examined all compounds containing fp2039, one of the promiscuous fingerprints ranked among the top three VIP features for multiple targets. We then examined CSNK1A1 degradation patterns induced by these compounds. Among compounds containing fp2039, we observed substantial heterogeneity in degradation outcomes; while a subset induced significant CSNK1A1 depletion (log_2_ fold-change < −0.415), others showed no activity despite containing the same key structural feature (Supplementary Fig. 19a). In line with this, our XGBoost model successfully predicted this context-dependent behavior, correctly distinguishing degraders from non-degraders among fp2039-containing compounds (Supplementary Fig. 19b).

To examine how individual fingerprints contribute to degradation predictions in different molecular contexts, we performed SHAP (SHapley Additive exPlanations) analysis on the ZFP91 model. SHAP assigns each input feature a contribution value for individual predictions, quantifying the positive or negative impact of each fingerprint on the predicted degradation. While VIP analysis identified fp1119 and fp47 among the most important features for ZFP91 degradation prediction (Fig. 4e), SHAP value distributions revealed substantial heterogeneity in how these fingerprints contributed to predictions across different compounds (Supplementary Fig. 20a). Although compounds containing fp1119 or fp47 showed a tendency toward stronger degradation, we identified a considerable overlap with non-degrading compounds (Supplementary Fig. 20b), confirming molecular context-dependent effects. To identify the molecular context governing fp1119-mediated degradation, we analyzed fingerprint co-occurrence patterns in ZFP91 degraders versus non-degraders containing fp1119. This analysis revealed several fingerprints significantly enriched in ZFP91-depleting compounds and others prevalent in inactive compounds. These results demonstrate that degradation outcomes are determined by combinations of structural features rather than isolated motifs (Supplementary Fig. 20c-d).

CSNK1A1 and WEE1 were degraded by 240 and 138 compounds, respectively, with an 85% co-occurrence rate. Given their high degradation frequency and the associated high model accuracy, we next investigated whether specific structural signatures governing their selectivity could be derived. Fingerprint enrichment analysis revealed distinct structural features driving WEE1 selectivity, with fp1698 emerging as the most discriminative motif between WEE1 and CSNK1A1 degradation (Fig. 4f and Supplementary Fig. 21a).

fp1698 encodes a direct carbon-carbon bond extending from the isoindolinone core, consistent with findings by Razumkov et al., who demonstrated that the nature of the carbon linkage at this position — specifically the presence or absence of a direct C−C bond — is a key determinant of WEE1 versus CK1α degradation selectivity^8^. Our machine learning approach recapitulated this finding, with compounds containing fp1698 showing selective WEE1 degradation in our experimental dataset (Supplementary Fig. 21b)

Beyond compound design, fingerprint-based models for frequently degraded neosubstrates could provide a practical tool to assess whether a compound of interest is likely to degrade common CRBN targets — enabling informed decisions about library enrichment or de-enrichment strategies. As an illustrative example, we selected neosubstrate–compound pairs from our screening data in which compounds degraded a given target without inducing significant degradation of CSNK1A1, WEE1, or ZFP91. Consistent with the observed selectivity, models predicted low degradation across all three frequent targets for these compounds, with predicted log₂ fold changes clustering near zero in contrast to the broader distribution observed across the full compound library (Supplementary Fig. 22).

## Discussion

Here, we report a large-scale proteomics-based screen of nearly 1,000 IMiD analogs to systematically map the CRBN neosubstrate target space. By employing MS-based quantification of global protein abundances, we determined the modulation of ∼10,000 endogenous proteins in response to MGD treatment. While earlier studies have established screening by proteomics as a gold standard tool for unbiased neosubstrate identification, our study extends these efforts by increasing scale, depth, throughput and robustness^6,22^. As advances in MS hardware continue to expand throughput, proteomics-based MGD drug screening is likely to become the standard approach in the future^51^.

To readily dissect CRL- and CRBN-dependent targets from independent regulations, we counter screened relevant hit compounds in presence of the NEDD8-activating enzyme inhibitor MLN4924 and in CRBN knockout cells. While most MGD-induced protein depletions were CRL- and CRBN-dependent, we identified a subset of recurring CRBN-independent regulations, including the endoplasmic reticulum (ER) resident proteins WLS and HMGCR. WLS is a multi-pass ER membrane protein that can be degraded by the ER-associated degradation (ERAD) pathway upon proteotoxic stress^52,53^. Its compound-induced depletion coincided with upregulation of several SREBP-1/2 target genes, including FASN, HMGCR and SQLE. This suggests a functional link between MGD-induced ER stress, and the induction of SREBP-1/2 target gene transcription^54^. Other than transcriptional control, HMGCR is also regulated post-translationally by sterols or non sterol isoprenoids levels^55^. Notably, two compounds induced HMGCR ubiquitination and depletion in a CRBN-independent manner, suggesting that they may directly or indirectly modulate sterol homeostasis. Together, these findings reveal previously undescribed off-target effects of CRBN-based MGDs that impact cholesterol metabolism.

We use global, site-specific analysis to profile MGD-induced ubiquitination events. Our analyses detected compound-induced ubiquitination for more than 230 proteins, including about 100 previously reported neosubstrates or candidate neosubstrates. These results not only validate our screening workflow but also underscore its power for *de novo* neosubstrate candidate identification. Both *in vitro* and cellular studies have previously demonstrated that protein ubiquitination in response to ubiquitin-proteasome system (UPS)-modulating drugs occurs within minutes^6,26,29^. Although ubiquitinomics alone cannot distinguish direct from indirect effects, we propose that well-characterized CRBN binders that induce ubiquitination of one or more sites within 30 minutes likely reflect direct degrader-neosubstrate relationships. To increase stringency, we integrated ubiquitinomics and global proteomics data, thereby further strengthening the nomination of high confidence neosubstrate candidates. Given that our screen was limited to a single cell line and primarily focused on IMiD derivatives, we have likely only uncovered only a small fraction of the potential neosubstrate space^56^. We anticipate that expanding screening efforts to include diverse chemical core structures and various biological models will significantly broaden this scope, potentially increasing the CRBN neosubstrate number to 1,000 or more proteins.

Structural G-loop motifs are the best characterized degrons for CRBN neosubstrates, with the majority of reported targets harboring such a sequence. However, the recent discovery of G-loop-independent neosubstrates, including VAV1, G3BP2, KDM4B and VCL, suggest a much broader target landscape^6,12–14^. Only about 50% of our newly discovered neosubstrate candidates contain a predicted G-loop degron, including the protein kinase IRAK1. Notably, our degrader exhibited high selectivity over IRAK2, 3 and 4, despite their high sequence homology, especially within the kinase domain. We mapped the IRAK1 interaction domain and identified the C-lobe of the kinase domain as a crucial determinant. Future studies, ideally guided by high-resolution protein structures are needed to precisely delineate the interaction interface and determine potential structural analogies with previously described G_-_loop-independent neosubstrates. The superior selectivity compared to traditional small-molecule inhibitors, combined with the ability to eliminate IRAK1’s non-catalytic scaffolding functions, demonstrates the therapeutic potential of the MGD modality.

BCL6 was previously identified as a CRBN-MGD neosubstrate through targeted zinc finger interactome and cellular reporter screens^16,57^. Western blot analysis and global proteomics demonstrated a weak, but significant downregulation of BCL6 by ALV1^16^. Our analyses identified three degrader probes that promoted both *in vitro* binding to CRBN and intracellular depletion of endogenous BCL6. For two of those, we additionally identified compound-specific intracellular ubiquitination. Remarkably, we were able to quantify BCL6 depletion, despite its very low expression level in HEK293 cells. AE-MS confirmed MGD-induced *in vitro* complex formation with CRBN and suggested that our compounds possess a higher affinity for the complex compared to ALV1. Reassuringly, this evidence translated to global proteomics, where our compounds induced a more robust depletion of BCL6 than ALV1. BCL6 contains three C_2_H_2_ zinc finger and three predicted G-loops. We provide experimental evidence that mutation of G608 alone is insufficient to abrogate the interaction with CRBN; however, a triple GN mutant disrupted the binding. These results are in line with recent findings suggesting that multiple zinc finger domains contribute to CRBN-MGD recruitment^17,49^.

Having generated proteome profiles for nearly 1,000 molecular glue degraders, we next utilized interpretable machine learning to systematically decode the structural determinants governing neosubstrate degradation. We selected XGBoost because gradient boosting models perform well on high-dimensional, sparse input features such as molecular fingerprints, while providing strong predictive accuracy and built-in feature importance measures. The observed variation in model performance across targets reflects differences in how predictable structure-activity relationships are for different neosubstrates. Some proteins may be more amenable to degradation by diverse chemical scaffolds, while others require specific structural features for productive ternary complex formation. The observed clustering patterns in VIP analysis and chemical space visualization support this interpretation. For example, ZFP91 and CSNK1A1 degraders showed clear separation, indicating that these two proteins have distinct structural requirements for degradation.

Our analysis showed that degradation outcomes emerge from combinations of structural features rather than from individual fingerprints alone. The presence of promiscuous fingerprints across multiple neosubstrates initially suggested that these features might be universal degradation determinants. However, several lines of evidence argue against this simple interpretation. First, analysis of promiscuous fp2039-containing compounds revealed some heterogeneity in CSNK1A1 degradation outcomes—while some compounds caused strong degradation, others showed no effect despite containing the same structural feature. Importantly, our models successfully predicted this context-dependent behavior, correctly distinguishing degraders from non-degraders among fp2039-containing compounds. Second, SHAP analysis revealed that individual fingerprints show context-dependent contributions—the same fingerprint can promote degradation in one molecular context while having minimal effect in another. Third, our co-occurrence analysis identified fingerprints that specifically enable or block degradation when present alongside promiscuous features. Together, these findings demonstrate that the molecular context surrounding a fingerprint determines its functional consequence, which is consistent with the known complexity of ternary complex formation where multiple structural elements must be present for productive degradation.

The network analysis revealed an interesting pattern: a densely connected module of fingerprints that frequently co-occur across top-ranked features for different genes. This module likely represents core structural elements necessary for CRBN engagement and ternary complex formation. In contrast, isolated fingerprints in the network appear to function as selectivity determinants. This organization suggests a hierarchical model where general CRBN-binding features provide the foundation, while additional structural features specify which neosubstrates get degraded.

Our validation through WEE1/CSNK1A1 selectivity analysis demonstrates that interpretable machine learning can successfully identify biologically relevant structural determinants. The identification of fp1698 as a WEE1 selectivity feature is particularly noteworthy because it was discovered through completely unbiased analysis of our large-scale proteomics data. The fact that this fingerprint corresponds to a specific chemistry previously shown to control selectivity by Razumkov et al. provides strong validation of our approach^58^.

While our compound library of approximately 1,000 MGDs demonstrates the applicability and utility of fingerprint-based machine learning for predicting neosubstrate degradation, the relatively modest library size limits model accuracy, particularly for less frequently degraded targets. Expanding the compound library would not only improve predictive performance for well-characterized neosubstrates but would also enable the development of reliable models for infrequently observed targets, which currently lack sufficient positive training examples. Future efforts combining larger and more structurally diverse compound libraries with the interpretable machine learning framework presented here are expected to substantially broaden the scope and accuracy of degradation prediction.

In summary, we present an MS-based screening and validation framework for the discovery of neosubstrates in an E3- and target-agnostic fashion. While we here focussed on CRBN-recruiting MGDs, our workflow is fully transferable to other E3 ligases and diverse chemical libraries. We envision proteomics becoming a standard screening tool in MGD drug discovery, offering a robust platform for both comprehensive neosubstrate identification and selectivity profiling.

## Methods

### Reagents

MLN4924, 2-chloroacetamide (CAA), tris(2-carboxyethyl)phosphine hydrochloride (TCEP), sodium deoxycholate, Na_2_HPO_4_, 3-(N-morpholino)propanesulfonic acid (MOPS), Tris(hydroxymethyl)-aminomethan, NP-40 alternative, glycerol, trifluoracetic acid, sodium chloride, formic acid and acetonitrile were from Merck. n-Undecyl-β-D-maltoside (UDM) from Anatrace. The MGD compound library was purchased from Enamine. Protease inhibitor mix from ThermoFisher Scientific (A32955). Trypsin from Promega. PTMScan® HS Ubiquitin/SUMO Remnant Motif (K-ε-GG) Kit (#59322) from Cell Signaling Technology.

### Cell culture, drug treatments and cell lysis (Proteomics)

HEK293 and HEK293T cells were from Cytion and were cultured in DMEM (VWR) supplemented with 10% FCS (Thermo Fisher Scientific). All compounds were dissolved in DMSO, to generate a 1000x stock solution. Compounds were plated in 96-well plates according to a randomized layout using an Opentrons OT-2. 40 (screening) or 27 (all follow-up experiments) compounds were allocated on each plate (in duplicate or triplicate, respectively), along with 16 or 15 DMSO controls, respectively. Cells were treated for the specified amount of time, washed with PBS and harvested using NEOsphere lysis buffer (0.05% UDM, 75 mM Tris-HCl pH 8.5, 40 mM CAA, 10 mM TCEP). The lysates were heated to 80°C for 10 min while shaking in a Thermomixer (Eppendorf). Proteins were digested by adding 400 ng of trypsin per sample (overnight, 37°C). The resulting peptides were desalted using in-house prepared, 200 µL two plug C18 StageTips (3M Empore^TM^)^59^ and then analysed by LC-MS/MS.

### Generation of stable cell lines

The plasmid vectors for stable cell line generation (overexpression of CRBN (CRBN_OE), CRBN knockout (KO) were from VectorBuilder. The guide RNA sequence for establishing CRBN-/- cells was as follows: ATCTAACTTCATGGCCTCGCGTTTTAGAGCTAGAAATAGCAAGTTAAAATAAGGCTAGTCCGTTATCAACTTGAAAAAGTGGCACCGAGTCGGTGC. HEK293 cells with stable gene overexpression or CRBN knockout were generated by lentiviral-mediated infection of cells according to published protocols^60^. In brief, HEK293T were co-transfected with psPAX2, pMD2.G, pRSV-Rev and a lentiviral construct for CRBN expression (CRBN_OE) or Cas9 and gRNA (CRBN_KO) under the control of a EF1A (CRBN_OE) or U6 (CRBN_KO) promoter. The supernatant was harvested after 48 hours of cell transfection and passed through a 0.45 µm filter. HEK293 cells were infected with the supernatant for 24 hours, followed by selection of transduced cells with puromycin (2 µg/ml). CRBN overexpression and knockout was verified by MS-based proteome profiling.

### Compound selection for follow-up experiments

All significantly downregulated proteins from the primary screen (adjusted *p*-value < 0.01) were extracted, and proteins significantly downregulated in the HEK293-GSPT1-G575N line were added. Highly active compounds (including GSPT1 degraders) were excluded from this selection. An additional filter requiring a π value (log_2_ (fold-change) x (-log_10_ adjusted p-value)) > 5 was applied. The corresponding compound-target pairs were were manually reviewed to remove likely false positives (i.e., those displaying large variance between treatment replicates or sparse identification based on a single peptide) and targets considered inaccessible to CRBN (e.g., secreted, mitochondrial, lysosomal or integral multi-pass membrane proteins). This led to the nomination of high confidence neosubstrate candidates across diverse functional classes. Finally, several positive controls (regulating previously reported neosubstrates) and selected highly promiscuous compounds (i.e., showing apparent simultaneous downregulation of many reported neosubstrates) were included in the final selection.

### Ubiquitinomics

Cell treatment and sample preparation was done according to our recently established protocol with a few modifications^26^. In brief, HEK293_CRBN_OE cells were cultured in 12-well plates and treated for 30 min with the specified compounds, followed by lysis with SDC buffer. Protein concentrations were determined using the BCA assay (Merck-Millipore) and the proteins were digested overnight at 37°C using 100:1 protein:trypsin ratio (Promega). After digestion, immunoprecipitation (IP) buffer (50 mM MOPS pH 7.2, 10 mM Na_2_HPO_4_, 50 mM NaCl) was added to the samples together with K-GG antibody-bead conjugate, followed by a 2 h incubation on a rotor wheel. Beads washing and peptide elution was performed according to manufacturer’s instructions. The peptide eluate was desalted using in-house prepared, 200 µl two plug C18 StageTips (3M EMPORE^TM^)^61^.

### Affinity enrichment mass spectrometry

The BCL6 (wt, G608N and 3xGN (G598N/G608N/G636N)) and IRAK1 (full-length, 1-381, 1-524) were from VectorBuilder. The respective constructs were transfected into HEK293 using PEI MAX (Polysciences #24765). The following day, the cells were resuspended in ice-cold NP-40 buffer (0.05% NP-40, 50 mM Tris-HCl pH 7.5, 150 mM NaCl, 5% glycerol), freshly supplemented with protease inhibitors. Cells were lysed by sonication followed by centrifugation at 20,000 ×g for 10 min (4 °C) and the supernatant transferred to a fresh tube. The protein concentration was determined using a BCA assay kit (Merck-Millipore) and the lysate concentration was adjusted to 1 mg/mL. Biotinylated CRBN-midi (CRELUX, WuXi AppTec) was added to the lysate along with DMSO/compound and incubated for 1 hour at 4 °C. Biotin affinity capture was used to isolate CRBN-glue-bound proteins (4 washes with lysis buffer after enrichment). Proteins were eluted with NEOsphere lysis buffer and digested by adding 100 ng of trypsin per sample (overnight, 37°C). The resulting peptides were desalted using in-house prepared, 200 µL two plug C18 StageTips (3M Empore^TM^)^59^ and then analysed by LC-MS/MS.

### LC-MS/MS measurements

Peptides were either analysed on mass spectrometers from Bruker (timsTOF pro2, timsTOF HT or timsTOF Ultra 2) or ThermoFisher (Orbitrap Astral). The LC Setup differed for the various sample types and/or mass spectrometers. For global proteomics on timsTOF instruments and global ubiquitinomics the following LC setup was used: Peptides were loaded on 30 cm reverse-phase columns (75 µm inner diameter, packed inhouse with ReproSil Saphir 100 C18 1.5 µm resin [Dr. Maisch GmbH]) using either a Vanquish™ Neo system (ThermoFisher) or a nanoElute® 2 system (Bruker). The column temperature was maintained at 60 °C using a column oven. The LC flow rate was 300 nL/min and the complete gradient was 50 minutes (global proteomics) or 45 minutes (ubiquitinomics). The LC setup of the samples measured with a higher throughput (global proteomics samples measured on the Orbitrap Astral instrument at ∼65 samples per day (SPD) or AE-MS samples measured on the timsTOF Ultra 2 at ∼100 SPD differed as follows: a 17 cm column with a 150 µm inner diameter was used, the flow rate was at 2,000 nL/min and the gradient length was 17 minutes (global proteomics) or 9 minutes (AE-MS), respectively. Data acquisition on timsTOF instruments was done using diaPASEF^62^ (proteomics) or slicePASEF^63^ (for ubiquitinomics). The diaPASEF acquisition scheme (ion mobility (IM) range from 0.65-1.35 and m/z range from 300-1,500) and was generated with py_diAID^64^. For global proteomics data measured on the Orbitrap Astral a Nanospray Flex™ source was used, with an ionization voltage of +2.2 kV, and an ion transfer tube temperature of 280 °C. The Orbitrap scan resolution was 240,000. The RF lens value was 40%, the AGC target 500%, and the maximum injection time 3 ms. A data-independent acquisition (DIA) scheme with 222 m/z windows of 3 to 10 Th covering an m/z range of 320-1231 was used, with the following Astral settings: scan range: 150 - 2,000 m/z, normalized HCD collision energy: 25%, AGC target: 800%, RF lens: 40%, maximum injection time: 3 ms.

### Raw data processing

MS raw files were analyzed using DIA-NN^65^ v2.1 enterprise (proteomics and AE-MS) or 2.2 enterprise (ubiquitinomics). Reviewed UniProt entries (human, SwissProt 01-2023 [9606]) were used as protein sequence database for DIA-NN searches. One missed cleavage, a maximum of one variable modification (Oxidation of methionines) and N-terminal excision of methionine were allowed. Carbamidomethylation of cysteines was set as fixed modification and K-GG (UniMod: 121) was added in case of ubiquitinomics. All data processings were carried out using library-free analysis mode in DIA-NN. --tims-scan was added as additional command in case of ubiquitinomics.

### Statistical data analysis

DIA-NN outputs were further processed with R. Peptide precursor quantifications with missing values in more than 50% of samples, or <75% of the DMSO-treated samples (for proteomics, <100% of compound-treated samples in case of ubiquitinomics) were discarded. Protein abundances were calculated using both precursor and fragment ion intensities (according to DIANN’s QuantUMS^66^ algorithm); K-GG peptide abundances were calculated based on precursor ion intensities levels using the MaxLFQ algorithm^67^, as implemented in the DIA-NN R package (https://github.com/vdemichev/diann-rpackage/). Complete missing cases in any of the conditions tested were rescued by accepting low-quality precursors (i.e., q-value > 0.01), where possible. For ubiquitinomics, missing values were imputed in DMSO in case of complete missingness. To minimize false positives, imputation was performed only when 100% condition completeness was achieved for a given peptide in at least one compound-treated condition. One ubiquitination site each mapping to SCRN3 and PRDM15 was quantified in only a single compound-treated condition; however, our imputation strategy failed to identify them as statistically significant. Due to their compound-specific induction, we nonetheless classified them as positive hits. K-GG peptide to site mapping was done using reviewed entries of the human UniProt database (SwissProt, release 01-2023). The protein (or peptide) intensities were normalized by median scaling and corrected for variance drift over time (if present) using the principal components (derived from principal component analysis (PCA)) belonging to DMSO samples. Subsequently, protein (or peptide, for ubiquitinomics) intensities were subjected to statistical testing with variance and log fold-change moderation using LIMMA^68^. p-values corrected for multiple hypothesis testing were used to assess significance for proteomics (q-value < 0.01) and ubiquitinomics (q-value < 0.05). For comparing proteome and ubiquitinome data, identifications were mapped at the gene level.

### Machine Learning

We trained XGBoost regression models to predict protein degradation from molecular fingerprints, performing 10-fold cross-validation with hyperparameter optimization via racing ANOVA. Chemical structures were represented as Morgan fingerprints (radius 2, 2,048 bits) using RDKit. For each gene, data were split into training (75%) and test (25%) with a fixed random seed to ensure reproducibility. Hyperparameters tuned included number of trees, features per split, minimum node size, and learning rate, with the best configuration selected based on cross-validation RMSE.

Model performance was evaluated on held-out test sets using Pearson correlation, R², and RMSE for regression. For binary classification, we applied a degradation threshold (log₂ FC < -0.41) and calculated accuracy, precision, recall, specificity, F1-score, and false positive rate from confusion matrices.

### Feature Importance and Interpretability

Variable Importance in Projection (VIP) scores quantified each fingerprint’s contribution to prediction accuracy. To assess stability, we trained models on each cross-validation fold and calculated mean and coefficient of variation of VIP scores across folds. Hierarchical clustering of VIP matrices identified genes with similar structural requirements.

SHAP (SHapley Additive exPlanations) analysis using TreeSHAP decomposed predictions into per-fingerprint contributions. SHAP value distributions across compounds revealed context-dependent effects of individual fingerprints.

Fingerprint co-occurrence and selectivity analyses used Fisher’s exact test with Benjamini-Hochberg correction to identify features enriched in specific compound categories. Important fingerprints were mapped to chemical structures using RDKit to highlight contributing atoms and bonds.

All analyses were performed in R using tidymodels and Python with RDKit. Statistical tests were two-sided with correction for multiple testing, where appropriate.

## Data availability

DIA-NN is freely available for academic use and can be downloaded at https://www.github.com/vdemichev/diann.

Source data are provided with this paper.

## Acknowledgements

Figures 1a and 4a were generated with Biorender (Agreement number RM27FBDPFP).

## Author contributions

Conceptualization and experiment design: H.D., M.S., B.Sh., B.Sch

Development of data analysis pipeline: B.Sch.

Data analysis and interpretation: H.D., M.S., B.Sh., D.W., P.Z., U.O.

Figure preparation: M.S., B.Sh., D.W.

Manuscript writing: M.S., B.Sh., H.D.

Proofreading and editing of the Manuscript: I.S., P.Z., U.O.

Development of NeosubstratesDB: I.S., B.Sh.

Execution of experiments: A.B., D.B., D.W., S.M., T.G.

All authors read and approved of the final manuscript.

## Author information

Correspondence and requests for materials should be addressed to

